# Lycopene Suppresses Lung Cancer Progression via PI3K/AKT Pathway Inhibition and Apoptosis Induction: Mechanistic and Safety Insights from Preclinical Models

**DOI:** 10.1101/2025.06.07.658457

**Authors:** Haibo Gu, Chengyu Pan, Qi Xu, Jingjing Lu, Tian Zhao, Keyan Fu, Xianting Yan, Yun Xu, Jian Ye

## Abstract

**Background:** Lung cancer remains a leading cause of cancer-related mortality, necessitating novel therapeutic strategies with minimal side effects. Lycopene, a natural carotenoid, has shown potential anticancer properties, yet its efficacy and mechanisms in lung cancer models require systematic exploration.

**Objective:** This study aimed to evaluate the antitumor effects of lycopene in both in vivo and in vitro lung cancer models, elucidate its molecular mechanisms, and assess systemic safety.

**Methods:** In vivo, tumor-bearing mice received intratumoral injections of lycopene at varying concentrations (10, 20, 40 mg/kg) to evaluate tumor regression . In vitro, A549 and LLC lung cancer cell lines were treated with lycopene (0–40 μM) to assess the ability of clonogenic survival, tumor new cell, cell invasion and promoting apoptosis.Western blot and immunohistochemistry were used to detect related pathways and apoptotic proteins, and target genes were silenced or overexpressed to verify the correctness of the pathways. Systemic toxicity was analyzed through blood biochemical profiling and histopathological examination (HE staining) of major organs at 40 mg/kg.

**Results:** Lycopene significantly reduced tumor volume in mice in a dose-dependent manner (p<0.05). Systemic toxicity assessments revealed no abnormalities in hepatic, renal, or hematological parameters, and organ histology remained unaffected at 40 mg/kg. In vitro, lycopene suppressed colony formation, tumor new cell, cell invasion numbers, and increased apoptosis rates about 60%. Mechanistically, lycopene downregulated phosphorylated PI3K and AKT levels, indicating pathway inhibition. Interference of PI3K gene silencing or overexpression suggests that the PI3K/AKT pathway is the main target for lycopene to produce apoptotic effects.

**Conclusion:** Lycopene exerts potent antitumor effects by inhibiting the PI3K/AKT pathway and promoting apoptosis in lung cancer cells, while demonstrating a favorable safety profile at therapeutic doses. These findings highlight its potential as an adjunctive therapy for lung cancer management.

## Introduction

Lung cancer remains one of the leading causes of cancer-related mortality worldwide, with non-small cell lung cancer (NSCLC) accounting for approximately 85% of cases [1, 2]. Despite advancements in targeted therapies and immunotherapies, challenges such as drug resistance and systemic toxicity persist, necessitating novel therapeutic strategies [3–5]. The PI3K/AKT signaling pathway, a critical regulator of cell survival, proliferation, and metastasis, is frequently dysregulated in lung cancer, making it a pivotal therapeutic target [6, 7]. Concurrently, natural compounds like lycopene — a carotenoid abundant in tomatoes—have garnered attention for their anticancer properties, particularly in modulating signaling pathways and oxidative stress [8–10]. This introduction synthesizes current knowledge on the role of PI3K/AKT in lung cancer progression and explores the potential of lycopene as a complementary therapeutic agent.

The PI3K/AKT pathway is activated by extracellular stimuli such as growth factors, leading to phosphorylation cascades that promote cell survival, angiogenesis, and epithelial-mesenchymal transition (EMT) [11–13]. In NSCLC, mutations in PI3K or AKT, or upstream regulators like EGFR, result in constitutive pathway activation, driving tumorigenesis and metastasis [14, 15]. For instance, phosphorylated AKT (p-AKT) enhances EMT by downregulating E-cadherin and upregulating N-cadherin and vimentin, facilitating cancer cell migration [16, 17]. Recent studies further highlight the spatial regulation of PI3K/AKT signaling within endosomal compartments, mediated by microtubule-associated proteins like MAP4, which influence tumor cell proliferation and invasion [18, 19]. However, existing PI3K inhibitors (such as alpelisib and copanlisib) face significant limitations in clinical application, on the one hand, their efficacy is highly dependent on the mutation status of PIK3CA and other genes, and the stratification of patients is complex, On the other hand, drug toxicity (e.g., hyperglycemia, neutropenia) limits long-term use [20–22]. Therefore, the exploration of natural compounds as novel regulators of PI3K/AKT pathway with low toxicity and broad spectrum effects has become a focus of current research.

Lycopene, a potent antioxidant, exhibits multifaceted anticancer effects. Epidemiological studies associate higher plasma lycopene levels with reduced lung cancer risk, particularly in smokers [23, 24]. Mechanistically, lycopene suppresses proliferation and metastasis by inhibiting matrix metalloproteinases (MMP-2/9), inducing cell cycle arrest at G0/G1 phase, and promoting apoptosis through modulation of Bcl-2/Bax and caspase-3 pathways [25, 26]. In vitro and in vivo studies demonstrate that lycopene reduces NSCLC cell viability and xenograft tumor growth while enhancing radiosensitivity [27, 28]. Emerging evidence suggests that natural compounds may enhance the efficacy of pathway-targeted therapies [29–31]. For example, traditional Chinese medicine formulations like Guben Jiedu Fang inhibit PI3K/AKT signaling, reversing EMT and reducing metastasis in NSCLC models [32]. Similarly, lycopene’s ability to downregulate MMPs and pro-survival proteins aligns with PI3K/AKT pathway modulation, though direct mechanistic links remain underexplored. Combining lycopene with PI3K/AKT inhibitors could potentially overcome resistance mechanisms while minimizing toxicity, leveraging both oxidative stress reduction and pathway suppression. However, at present, no lycopene has been reported to have an effect on lung cancer through its effect on PI3K signaling pathway. Therefore, the main purpose of this paper is to explore whether lycopene can ultimately achieve therapeutic purposes through its effect on PI3K signaling pathway. It provides a new theoretical basis for further clinical administration alone or combined therapy.

## Materials and methods

### Animals and groups

All mice studies were conducted by the Zhejiang University Animal Study Committee requirements for the care and use of laboratory animals in research(No.ZJU20241025). 6 weeks old female Balb/c nude mice were obtained from Shanghai Bioscience Co. Ltd (Shanghai, China). All mice were housed under standard conditions with a humidity of 50% and a temperature of 20–22°C, following a 12-h light/dark cycle. They were provided with ad libitum access to water and standard rodent chow.

Cells in the logarithmic growth phase were trypsinized and centrifuged at 1000 × g for 5 min. After washing with phosphate-buffered saline (PBS), the cells were resuspended in PBS to prepare a single-cell suspension at a density of 10^7^ cells/mL. A 50 μL aliquot of the suspension was subcutaneously inoculated into the right axilla of nude mice. Two weeks post-inoculation, when tumors had enlarged, 32 mice were randomly allocated into four groups (n = 8/group): (1) control group; (2) low-dose lycopene group; (3) medium-dose lycopene group; and (4) high-dose lycopene group. The control group received intratumoral injections of 10 μL saline, while the lycopene-treated groups were administered 10 μL lycopene solutions at doses of 10, 20, or 40 mg/kg, respectively. All injections were performed every other day. On day 20 post-treatment, all mice were humanely euthanized via cervical dislocation. Tumor volumes were measured, and the tumor inhibition rate was calculated using the formula:Tumor inhibition rate (%) = [(Mean tumor weight of control group − Mean tumor weight of treatment group) / Mean tumor weight of control group] × 100%.

Immediately after excision, tumor tissues were bisected. One portion was fixed in 10% neutral-buffered formalin for routine paraffin embedding, followed by hematoxylin-eosin (HE) staining and immunohistochemical analysis. The other portion was snap-frozen in liquid nitrogen for subsequent protein extraction and evaluation of tumor-associated protein expression.

### Hematoxylin-eosin (HE) staining

Histological analysis was performed as follows: the isolated tumor tissues were fixed in 4% paraformaldehyde solution, embedded in paraffin, and subsequently sectioned into 6-μm-thick slices using a microtome (Leica, Biocut, Germany). The sections were stained with hematoxylin and eosin (H&E) in accordance with the manufacturer’s protocol.

### Immunohistochemical (IHC) staining

The tumor sections (6-μm) were deparaffinized using xylene-ethanol gradient hydration, followed by antigen retrieval. Endogenous peroxidase activity was subsequently blocked with 5% hydrogen peroxide. After 30-minute blocking with 5% BSA, the sections were incubated overnight at 4°C with primary antibodies against Ki67, p-PI3K, p-AKT and cleaved-caspase-3. The following day, the sections were sequentially treated with anti-rabbit immunoglobulin and streptavidin-conjugated horseradish peroxidase (HRP). Final detection was achieved using 3,3’-diaminobenzidine (DAB) chromogenic substrate, with hematoxylin counterstaining applied for nuclear visualization.Photographs were captured using a microscope (3DHISTECH Pannoramic SCAN II, Hungary). Brown staining in cytoplasm and nucleus were considered as positive. And the ratios of brown staining areas to total areas were calculated blindly.

### Biochemical assays

The blood samples were incubated at 4°C for 4 hours, followed by centrifugation at 3000 × g for 20 min under 4°C to separate the serum. The obtained serum was subsequently analyzed for aspartate aminotransferase (AST), alanine aminotransferase (ALT), blood urea nitrogen (BUN), and creatinine (CREA) using a serum biochemical analyzer (FUJI DRI-CHEM NX700i, USA) in accordance with the manufacturer’s protocols. All measurements were documented systematically during the analytical process.

### Western blotting

Proteins extracted from tissues or cells were separated by sodium dodecyl sulfate-polyacrylamide gel electrophoresis (SDS-PAGE) and subsequently transferred onto polyvinylidene difluoride (PVDF) membranes. The membranes were probed with antibodies against p-PI3K((#4228), PI3K(20584-1-AP), p-AKT(66444-1-Ig), AKT(10176-2-AP), Bcl-2(#3498), Bax(#2772) and β-actin(66009-1-Ig) for immunodetection. Protein quantification was performed using Image Lab software (Bio-Rad, Chemidoc MP, USA) with optical density units, and the results were normalized to the expression levels of β-actin(66009-1-Ig) in corresponding samples.

### Cell Culture

BEAS-2B, A549 and LLC cells were cultured in DMEM medium supplemented with 10% fetal bovine serum (FBS), 50 U/ml penicillin, and 50 U/ml streptomycin, and maintained at 37°C in a humidified atmosphere containing 5% CO₂. Upon reaching 80% confluence, the cells were subcultured or transferred to designated culture plates for subsequent experimental procedures.

### Cell viability assay

BEAS-2B, A549 and LLC cells were seeded in 96-well plates (2 × 10⁴ cells/well in 100 μL medium) and allowed to adhere overnight. Following complete cell attachment, the cells were treated with gradient dilutions of lycopene for 24 h. Subsequently, CCK-8 reagent dissolved in serum-free DMEM (adjusted to 10% working concentration) was added (100 μL/well) and incubated for 1 h. Optical density was then measured at 450 nm using a microplate reader(Tecan spark, USA).

### Cell wound scratch assay

The cell scratch assay was performed to evaluate cell migration ability. A549 and LLC cells were seeded into 6-well plates respectively. After reaching complete confluence, straight scratches were created in the cell monolayer using a sterile 10 μL pipette tip. The cells were then treated with lycopene at concentrations of 0, 10, 20, and 40 μM in DMEM supplemented with 1% (FBS) for 48 hours. Cell migration distance was quantitatively assessed by measuring the wound width at 0 h and 48 hours post-treatment using microscopic image analysis.

### Clone formation assay

Cells were seeded in 6-well plates at a density of 1,000 cells per well. Following respective treatments, the cells were maintained in culture for 14 days, with medium replacement performed every 3 days. After the 14-day incubation period, cells were fixed with methanol and stained with 0.2% crystal violet solution. Colony formation was subsequently documented through microscopic photography and quantitatively analyzed by manual counting of stained colonies.

### EDU assay

Cells in logarithmic growth phase (2×10⁴ cells/well) were seeded into 96-well plates and cultured overnight. Subsequently, the cells were incubated with various concentrations of lycopene for 24 h. Following treatment, cells were washed and fixed with methanol for 30 minutes. After two PBS washes, cell permeabilization was performed using 0.3% Triton X-100 in PBS, followed by Edu reaction solution staining according to the manufacturer’s protocol. Fluorescence microscopy imaging was conducted to capture cellular morphology and staining patterns.

### Flow cytometry

Cells (5 × 10⁵ cells/well) were seeded in 6-well plates and cultured overnight in complete DMEM medium. Subsequently, the medium was aspirated and replaced with either 2 mL fresh complete DMEM medium (control group) or equivalent volume of lycopene solutions prepared in complete DMEM medium (final concentrations: 10, 20, and 40 μM) for the lycopene-treated groups. Following 24 h of additional incubation, cells were harvested and subjected to apoptosis analysis using flow cytometry. Briefly, 5 μL Annexin V-FITC and 10 μL propidium iodide (PI) staining solutions were sequentially added to each sample, followed by 15 min of dark incubation prior to detection.

### Immunofluorescence staining

Cells treated with varying concentrations of lycopene were fixed with methanol for 15 min, followed by blocking with 5% BSA at room temperature for 30 min. After blocking, cells were incubated overnight at 4°C with the primary p-PI3K antibody. On the following day, samples were treated with Alexa fluor 488 anti-rabbit secondary IgG antibody (Thermo Fisher, USA) at room temperature for 1 h. Nuclei were counterstained with DAPI, and fluorescent signals were subsequently captured and analyzed using laser scanning confocal microscopy(Olympus SpinS R, Japan).

### Cell transfection

The experimental protocol was conducted as follows: A549 and LLC cells were transfected with PI3K-specific small interfering RNA (siRNA) or PI3K cDNA plasmid. Cells were seeded at a density of 1×10⁵ cells per well in 6-well plates and cultured for 24 hours. Following cell adhesion, PI3K-specific siRNA or cDNA-PI3K plasmid was carefully combined with Lipofectamine 3000 reagent according to manufacturer’s instructions. The transfection complexes were gently mixed and then gradually added to the cell culture medium through dropwise administration. After gentle agitation to ensure uniform distribution, transfected cells were maintained under standard culture conditions for 6 hours. Subsequently, the transfected cells were exposed to lycopene (40 μM) for 24 hours. Finally, cells were harvested and lysed for subsequent Western blot analysis to evaluate protein expression levels.

### Statistical analysis

The data are expressed as mean ± standard deviation (SD). The differences between the two groups were analyzed using unpaired t-test. A p-value < 0.05 was considered statistically significant. All statistical analyses were performed with GraphPad Prism 8.0 (GraphPad Software).

## Results

### Lycopene reduces the growth of tumors

As shown in Fig. 1a and b, the lycopene-treated groups exhibited a significant dose-dependent reduction in tumor volume compared with the model group. Tumor inhibition rates reached 34.6%, 60.9%, and 85.2% in the low-, medium-, and high-dose lycopene groups, respectively (P < 0.05, Fig. 1c). Corresponding hematoxylin and eosin (H&E) staining results in Fig. 1d revealed varying degrees of necrotic areas within tumors from lycopene-treated groups. Furthermore, immunohistochemical analysis demonstrated that lycopene treatment markedly suppressed the expression of the proliferation marker Ki-67 in tumor tissues in a concentration-dependent manner (Fig. 1e). These findings collectively indicate that lycopene significantly inhibits tumor growth in tumor-bearing nude mice.

**Figure 1.**
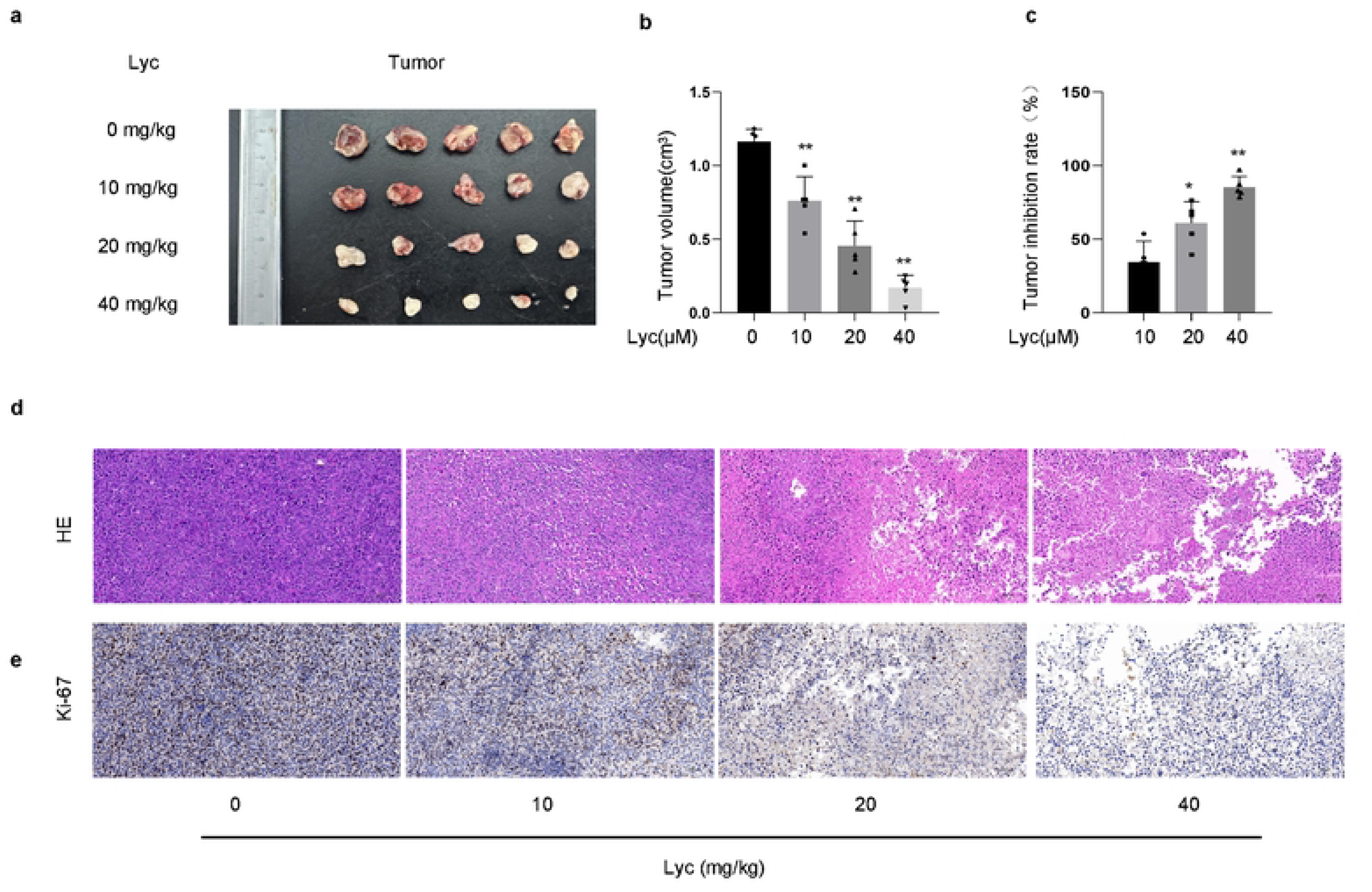
Lycopene suppresses tumor growth in nude mouse xenograft models. (a, b) Tumor volume measurements showing dose-dependent inhibition of tumor growth in lycopene-treated groups compared to the model group. (c) Tumor inhibition rates for lycopene groups, respectively. (d) Hematoxylin and eosin (H&E) staining of tumor tissues. (e) Immunohistochemical analysis of Ki-67 expression. Values are expressed as mean ± SD (n ≥4). *P<0.05, **P < 0.01 vs control group.

### Lycopene promotes mitochondrial apoptosis and related proteins on tissue

The Bax/Bcl-2 ratio serves as a pivotal determinant in regulating cellular susceptibility to apoptosis suppression. Modulation of their expression levels or interaction may regulate the apoptotic process. As illustrated in Figure 2a, lycopene administration significantly downregulated Bcl-2 expression and upregulated Bax protein levels in tumor tissues compared with the model group, with these effects exhibiting dose-dependent enhancement. This dose-responsive pattern suggests lycopene may promote mitochondrial-mediated apoptosis in tumor cells. Furthermore, phosphorylated PI3K (p-PI3K) and AKT (p-AKT) exhibited marked decreases in expression with increasing lycopene concentrations, while their respective total protein levels remained unaltered. These findings imply that lycopene potentially modulates tumor cell apoptosis through inhibition of p-PI3K signaling pathways.

**Figure 2.**
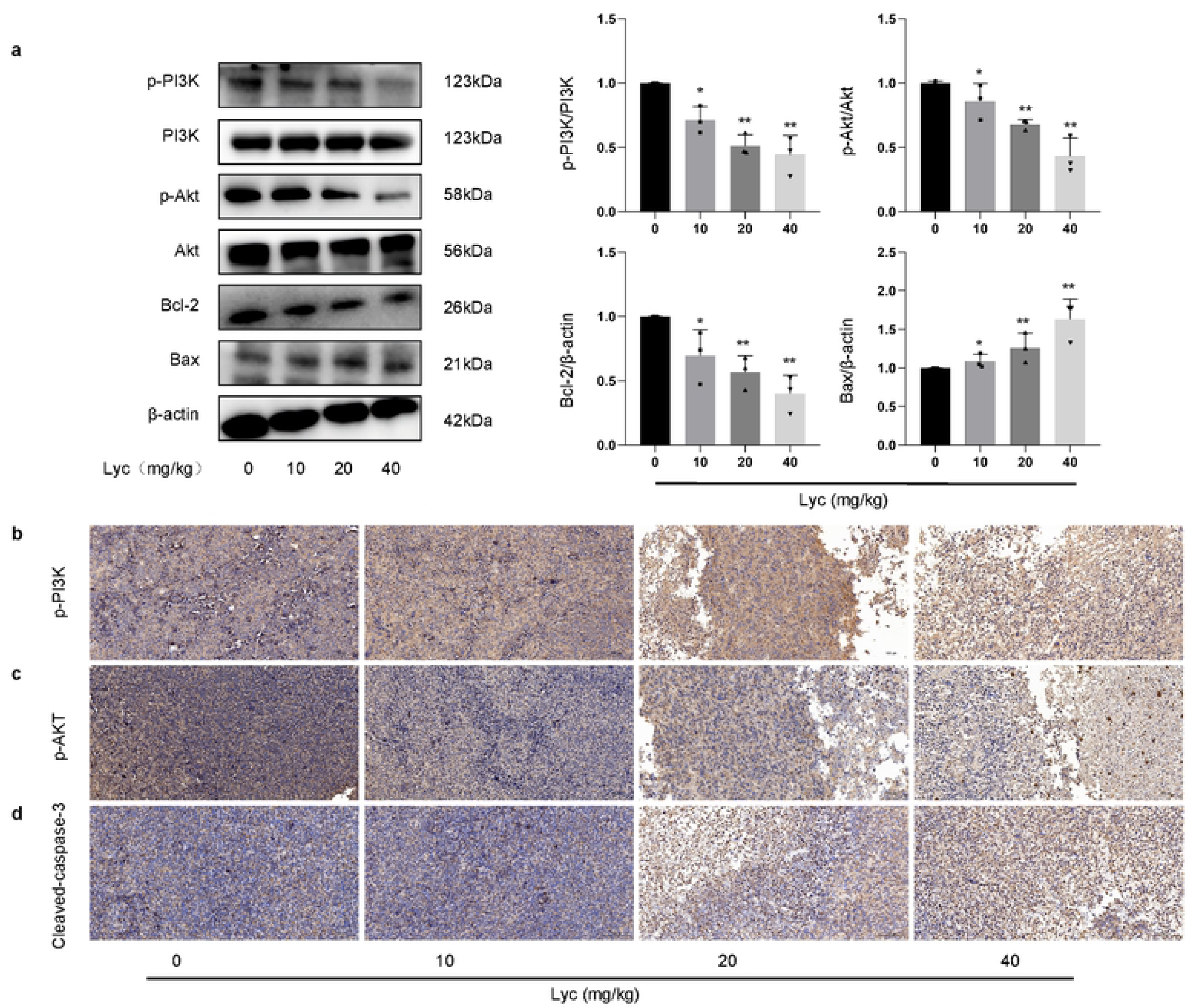
Lycopene modulates mitochondrial apoptosis and PI3K/AKT signaling in tumor tissues. (a) Western blot analysis of Bax, Bcl-2, p-PI3K, and p-AKT in tumor tissues from lycopene-treated tumor-bearing mice. (b-d) Immunohistochemical staining of p-PI3K, p-AKT, and cleaved caspase-3 in tumor tissues. Values are expressed as mean ± SD (n ≥3). *P<0.05, **P < 0.01 vs control group.

Immunohistochemical staining was subsequently conducted to examine the expression patterns of two critical signaling regulators, p-PI3K and p-Akt. The analytical results presented in Figure 2b, c demonstrate that lycopene administration in tumor-bearing mice significantly attenuated the immunopositive areas of these biomarkers. Notably, this suppressive effect exhibited a dose-dependent response, with higher lycopene concentrations correlating with more substantial reductions in positive staining areas.At the same time, the positive area of Cleaved-caspase-3 increased significantly with the increase of lycopene concentration showed in Figure 2d.

### Biocompatibility of lycopene

To evaluate the biosafety of lycopene, mice were intraperitoneally administered a high-concentration lycopene solution (40 mg/kg) for physiological and pathological assessment. Histopathological examination on day 10 post-injection revealed no significant pathological alterations in major organs (Figure 3a). Furthermore, long-term hematological toxicity monitoring demonstrated absence of systemic toxicity, as evidenced by statistically insignificant variations in serum biomarkers (ALT, AST, BUN, and CREA) across different time intervals showed in Figure 3b. These findings collectively indicate that the lycopene formulation possesses favorable biocompatibility.

**Figure 3.**
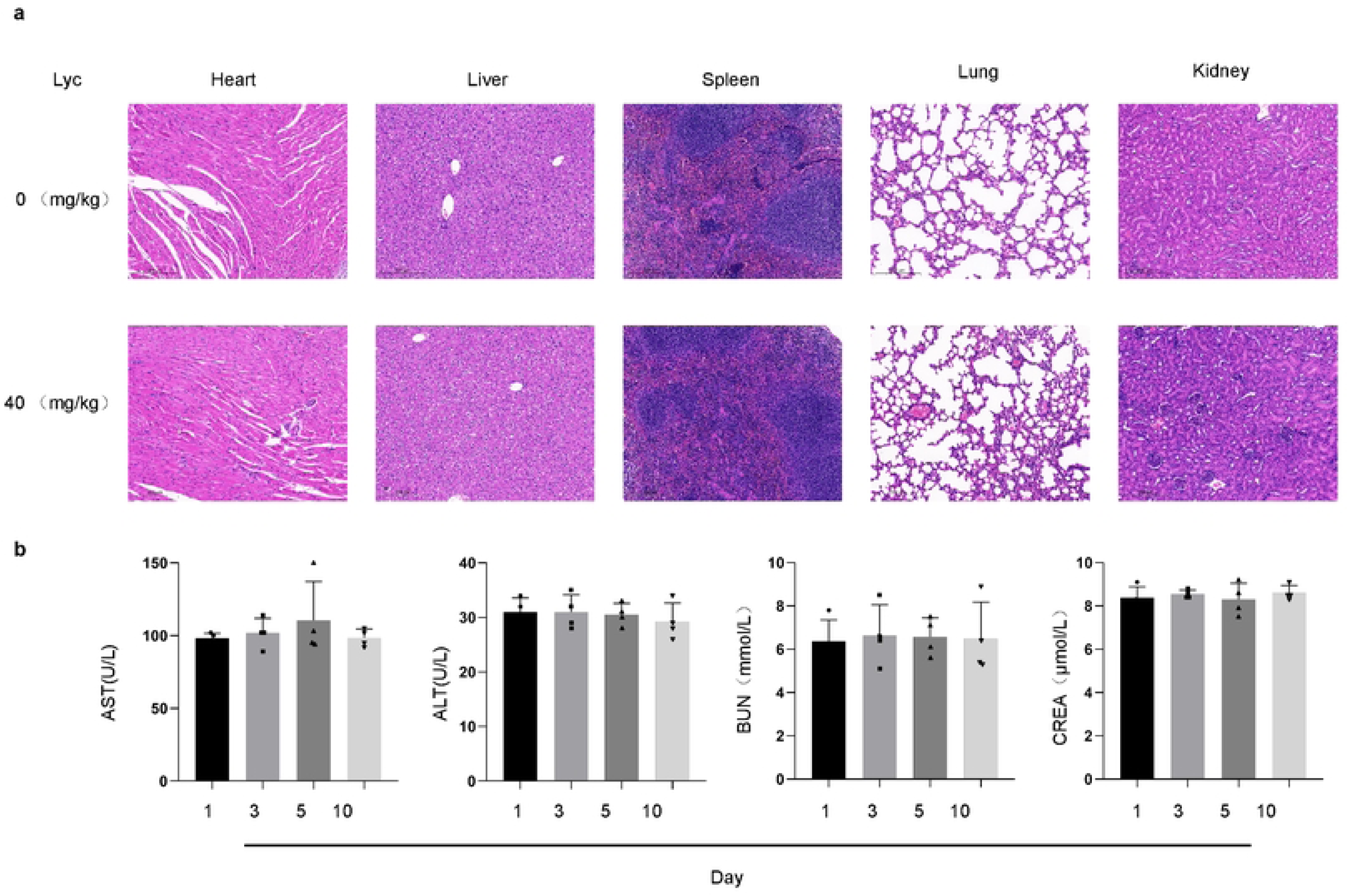
Evaluation of lycopene biosafety and systemic biocompatibility. (a) Histopathological analysis of major organs (heart, liver, spleen, lung, and kidney) in mice intraperitoneally administered a high-concentration lycopene solution (40 mg/kg) for 10 days. (b) Long-term hematological toxicity assessment of lycopene for serum biomarkers, including ALT, AST, BUN, and CREA. Values are expressed as mean ± SD (n ≥4). *P<0.05, **P < 0.01 vs control group.

### Tumor cell killing of lycopeneon cell viability

To further investigate the mechanism underlying lycopene-mediated attenuation of lung cancer progression in mice, we conducted in vitro experiments utilizing four cellular models: BEAS-2B (normal bronchial epithelial cells), PC9, A549, and LLC (lung cancer cell lines). As demonstrated in Figure 4a, BEAS-2B normal lung cells exhibited cytotoxic responses only when exposed to lycopene concentrations reaching 160 μM. In contrast, the lung cancer cell lines (PC9, A549, and LLC) manifested significant cytotoxicity at substantially lower concentrations (10-20 μM), with a dose-dependent escalation of toxic effects observed following increased lycopene administration. These findings collectively suggest that pulmonary tumor cells demonstrate enhanced sensitivity to lycopene-induced cytotoxicity compared to normal lung cells.

**Figure 4.**
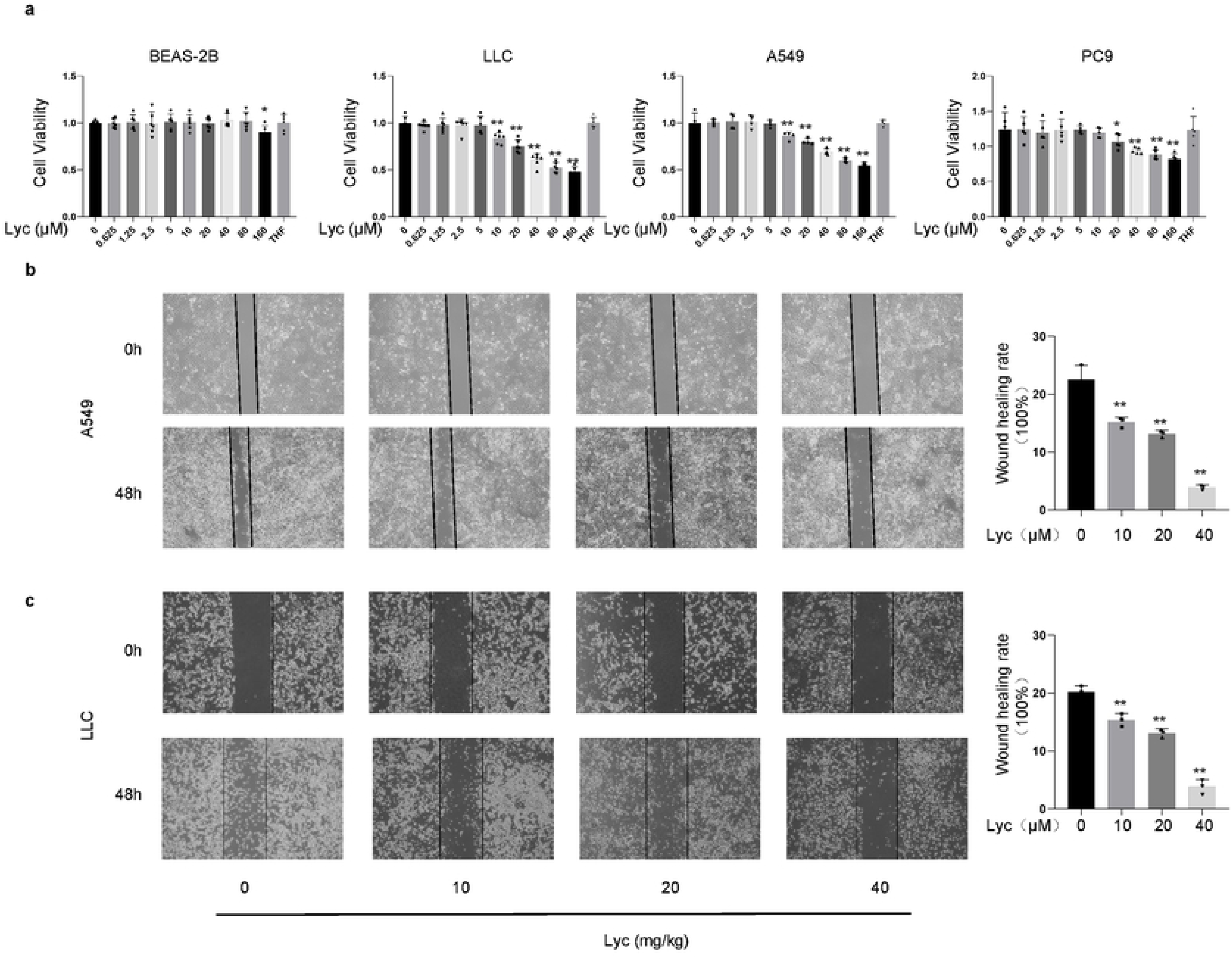
Lycopene selectively inhibits lung cancer cell viability and migration in vitro. (a) Cytotoxicity assessment of lycopene in normal bronchial epithelial cells (BEAS-2B) and lung cancer cell lines (PC9, A549, LLC). (b, c) Scratch assay analysis of lycopene effects on A549 and LLC cell migration. Values are expressed as mean ± SD (n ≥3). *P<0.05, **P < 0.01 vs control group.

Figures 4b and 4c present the scratch assay results of A549 and LLC cells, respectively. The results demonstrated that lycopene treatment significantly inhibited cell migration after 48 hours of exposure. At a concentration of 40 mg/kg, the migration rates of A549 and LLC cells were reduced to 3.9% and 3.8% respectively after 48-hour treatment, in contrast to the untreated groups which exhibited substantially higher migration rates of 22.6% and 20.2% for A549 and LLC cells, correspondingly.

### Enhanced tumor cell migration, invasion and proliferation of lycopene

The cell proliferation rate was quantitatively assessed using the EdU labeling assay to measure stage-specific newly generated cells. As demonstrated in Figures 5 a,b,c,d, the results revealed that within 24 hours, the lycopene-treated groups exhibited a significant reduction in newly generated cells compared to untreated controls in both A549 and LLC cell lines. Specifically, the untreated groups displayed 17.5 and 18.6 new cells for A549 and LLC, respectively, whereas administration of 40 mg/kg lycopene markedly decreased these values to 3.9 and 5.4 new cells, corresponding to a 77.7% and 70.9% inhibition rate relative to controls. The cell colony formation assay demonstrated that lycopene dose-dependently inhibited the proliferation of lung cancer cells in Figures 5e,f,g,h.

**Figure 5.**
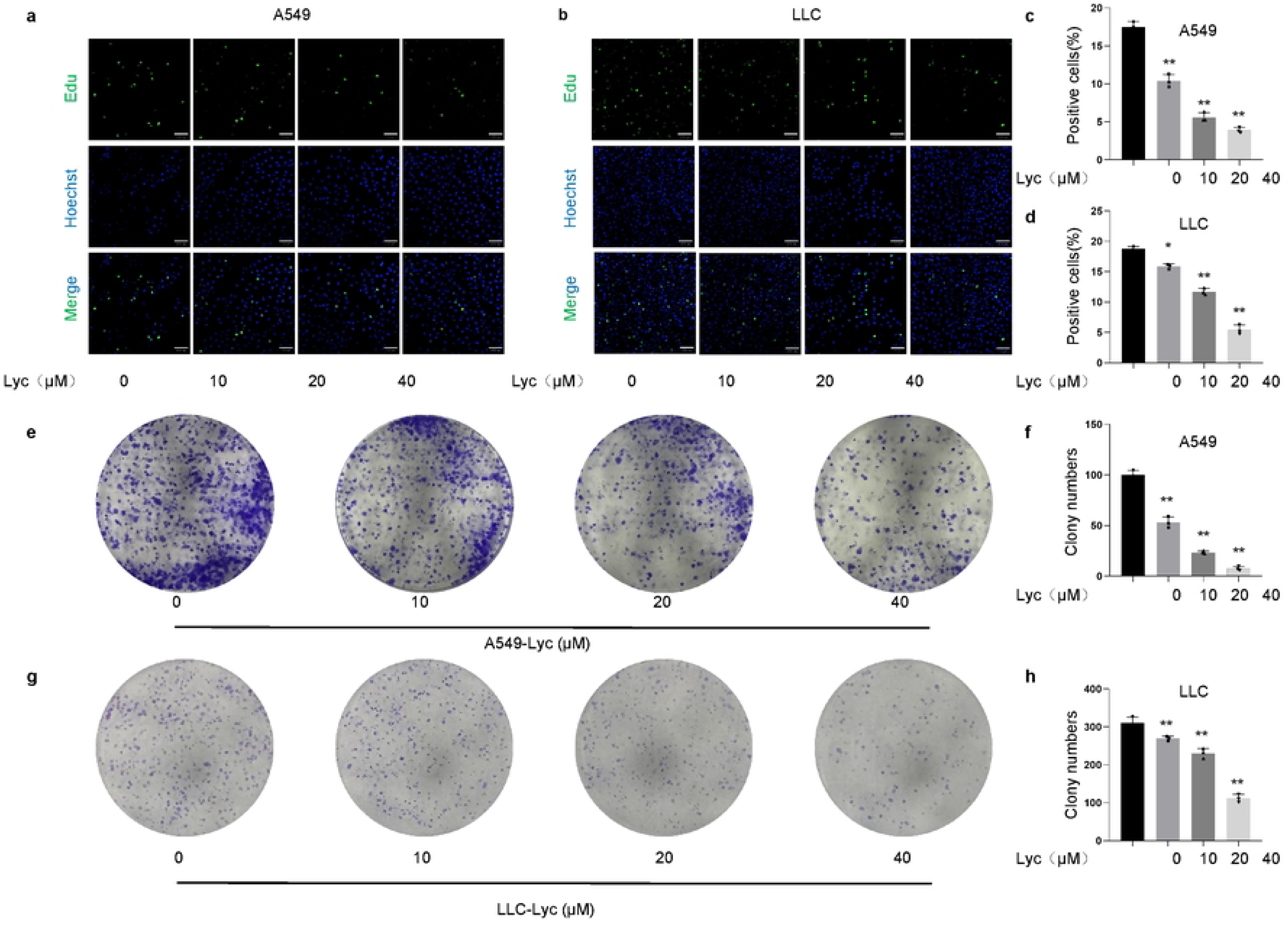
Lycopene suppresses proliferation and colony formation of lung cancer cells in vitro. (a-d) EdU labeling assay quantifying newly generated A549 and LLC cells after 24-hour lycopene treatment. (e-h) Colony formation assay revealing dose-dependent inhibition of lung cancer cell proliferation by lycopene. Values are expressed as mean ± SD (n = 3). *P<0.05, **P < 0.01 vs control group.

### Lycopene promotes the apoptosis of lung cancer cells and inhibits their invasion

As shown in Figure 6a and 6b, treatment with lycopene at doses of 10, 20, and 40 mg/kg significantly increased the apoptosis rates of A549 and LLC cells, suggesting that lycopene may effectively promote tumor cell apoptosis. As depicted in Figure 6c and 6d, lycopene treatment significantly suppressed the invasive capacity of A549 and LLC cells in a concentration-dependent manner. Transwell assay revealed that 40 μM lycopene treatment for 24 hours reduced the number of invaded cells to 59.33 and 74.12 for A549 and LLC cells, respectively, compared with 26,265 and 33,123 cells in the untreated control group. Notably, statistically significant differences in inhibitory effects were observed between all concentration groups (10 μM, 20 μM, and 40 μM) and the untreated control group.

**Figure 6.**
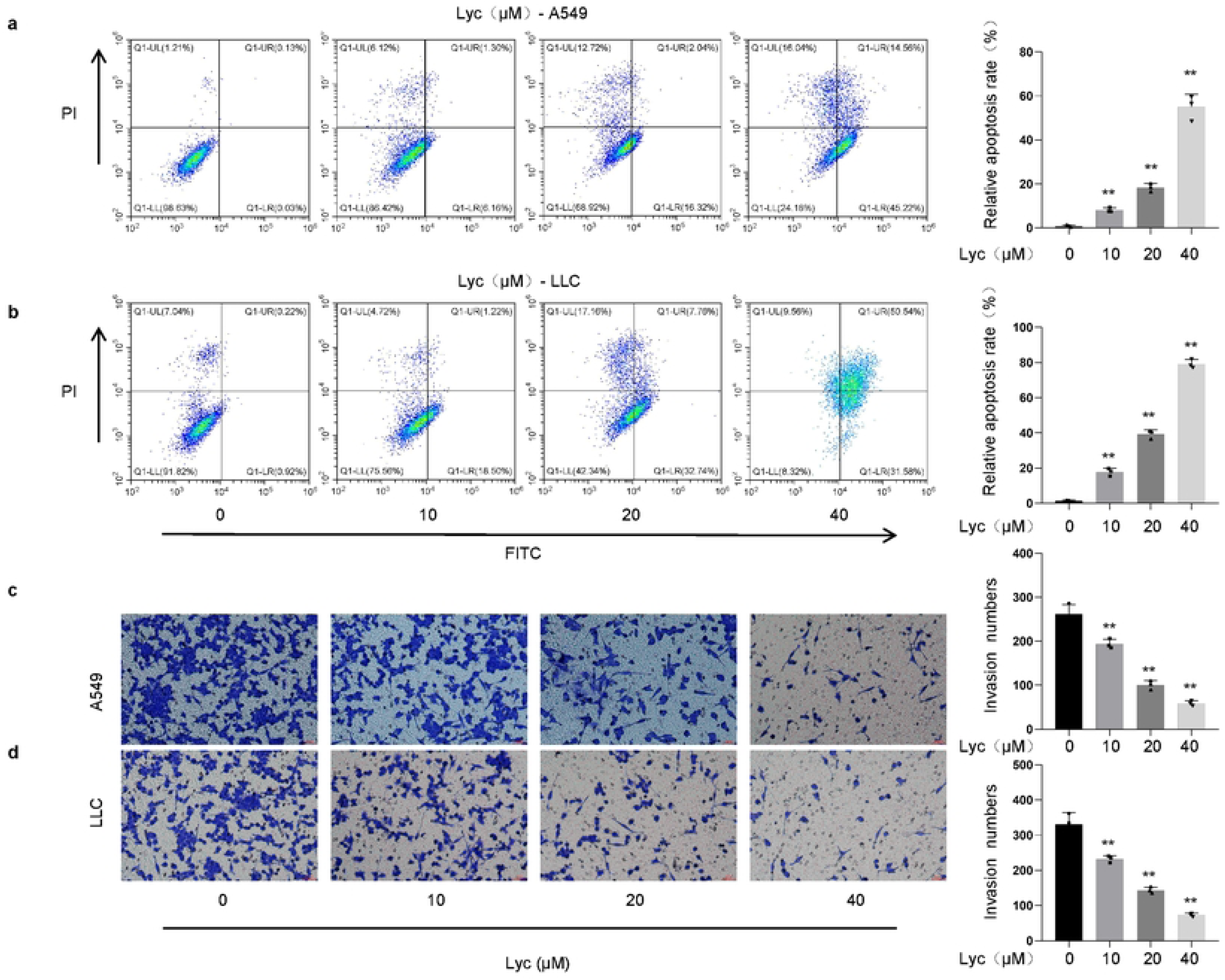
Lycopene induces apoptosis and suppresses invasion in lung cancer cells. (a, b) Apoptosis rates of A549 and LLC cells following lycopene treatment (10, 20, 40 mg/kg). (c, d) Transwell invasion assay demonstrating lycopene’s concentration-dependent inhibition of A549 and LLC cell invasiveness. Values are expressed as mean ± SD (n = 3). *P<0.05, **P < 0.01 vs control group.

### Role of PI3K/AKT signaling pathway in lycopene-exerted Tumor cell killing

The results presented in Figures 7a and 7b demonstrate that hyperphosphorylation of PI3K was observed in both A549 and LLC cells. Lycopene pretreatment (10–40 μM) significantly reduced the phosphorylation levels of iPI3K in a concentration-dependent manner. Furthermore, lycopene (10–40 μM) pretreatment markedly downregulated the expression of p-AKT and Bax proteins while upregulating Bcl-2 protein expression. Notably, lycopene exhibited no significant effect on the total protein levels of PI3K and AKT. Collectively, these findings suggest that lycopene may induce apoptosis by inhibiting the phosphorylation of PI3K, thereby suppressing AKT activation and subsequently modulating Bcl-2 family protein expression.

**Figure 7.**
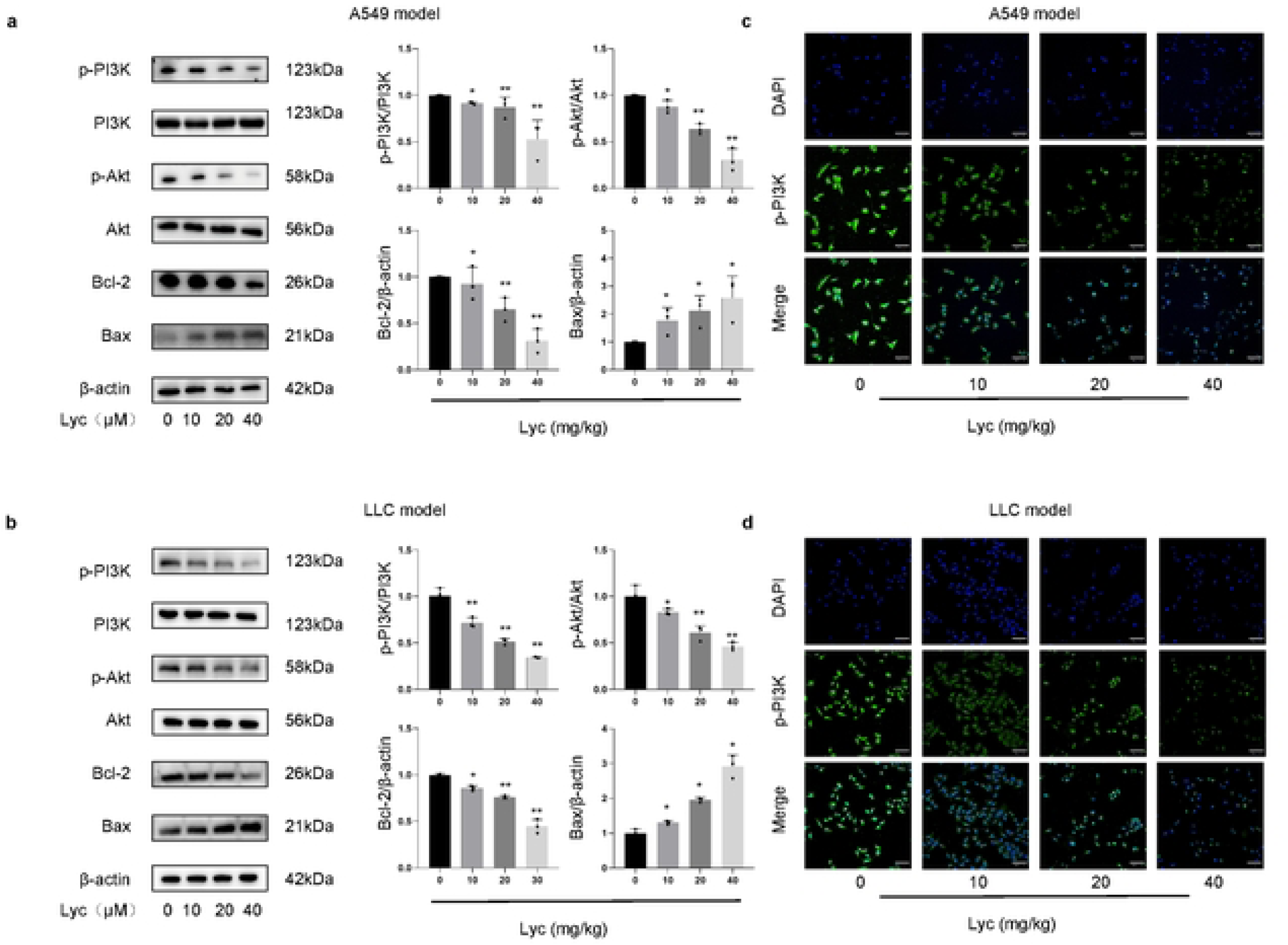
Lycopene inhibits PI3K/AKT signaling and modulates apoptotic protein expression in lung cancer cells. (a-b) Western blot analysis of p-PI3K, p-AKT, Bax, and Bcl-2 in A549 and LLC cells treated with lycopene (10–40 μM). (c, d) Immunofluorescence analysis of p-PI3K expression in A549 and LLC cells. Values are expressed as mean ± SD (n = 3). *P<0.05, **P < 0.01 vs control group.

We also performed immunofluorescence analysis to detect the expression of p-PI3K, as shown in Figs. 7c and d. The fluorescence intensity of p-PI3K in both cell types gradually decreased with increasing concentrations of lycopene, demonstrating consistent trends with the Western blot results. Furthermore, the examination of the entire cell signaling pathway corroborated our findings from the animal experiments.

### Lycopene suppresses PI3K signaling to induce apoptosis in A549 and LLC cells

To assess the correlation between lycopene-induced regulation and PI3K suppression, A549 and LLC cells transfected with PI3K siRNA or cDNA were utilized. Our findings demonstrated that PI3K siRNA transfection significantly reduced the expression of PI3K, p-PI3K, and other apoptosis-related proteins, including the p-AKT and Bax/Bcl-2 ratio. These outcomes partially mimicked the lycopene-induced effects. Notably, combined treatment with PI3K siRNA transfection and lycopene failed to induce further reductions in p-PI3K levels or apoptosis-associated proteins (Figures 8a, b). Conversely, PI3K cDNA transfection markedly enhanced PI3K expression, which was accompanied by elevated p-AKT and Bcl-2 levels. Lycopene pretreatment neither reversed the downregulated p-PI3K nor attenuated the associated p-AKT and Bax proteins (Figures 8c, d). These collective findings suggest that the pro-apoptotic effect of lycopene in tumor cells is likely dependent on its inhibitory effect on p-PI3K signaling.

**Figure 8.**
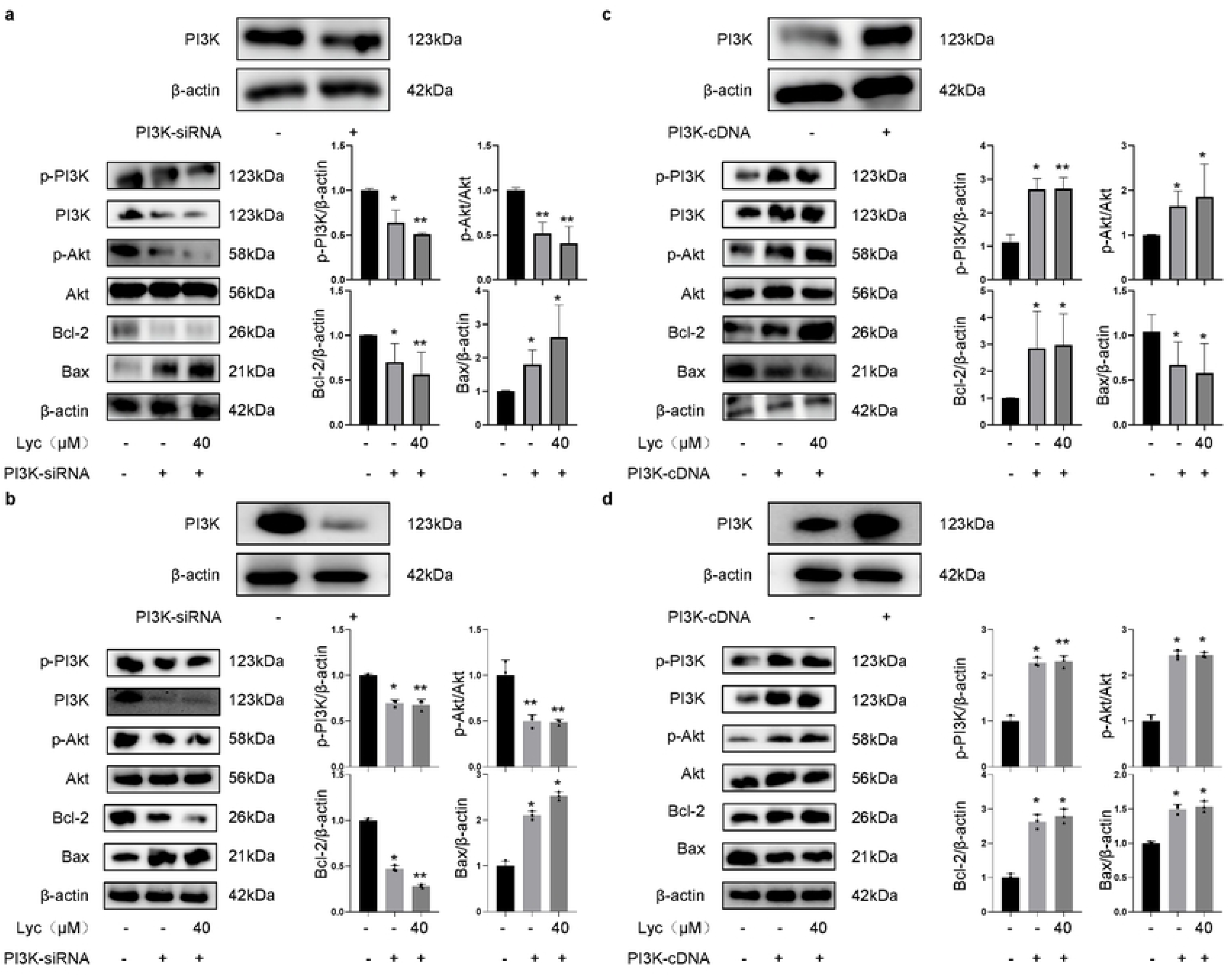
Lycopene suppresses PI3K signaling resulting in the induction of apoptosis in A549 and LLC cells. (a, b) Western blot analysis of PI3K, p-PI3K, p-AKT, AKT, Bax, and Bcl-2 in cells transfected with PI3K siRNA or treated with lycopene. (c, d) Overexpression of PI3K via cDNA transfection elevated PI3K, p-PI3K, p-AKT, AKT, Bax, and Bcl-2 levels.Values are expressed as mean ± SD (n = 3). *P<0.05, **P < 0.01 vs control group.

## Discussion

The present study systematically investigated the anti-tumor effects and molecular mechanisms of lycopene in both in vivo and in vitro models, revealing its dose-dependent tumor suppression, non-toxic safety profile, and potential signaling pathway regulation.This dual advantage of efficacy and safety addresses a critical challenge in current chemotherapeutic strategies, where dose-limiting toxicity often compromises therapeutic outcomes. These findings not only align with previous studies on lycopene’s anti-cancer properties but also provide novel insights into its mechanism of action in lung cancer models.

Our in vivo experiments demonstrated that intratumoral injection of lycopene (10–40 mg/kg) significantly reduced tumor volume in a dose-dependent manner, corroborating earlier reports on lycopene’s anti-tumor efficacy in breast and prostate cancers [9, 33]. And the intratumoral administration of lycopene (10-40 mg/kg) induces dose-dependent tumor regression in murine models, aligning with previous reports on carotenoids’ anti-cancer potential [34]. Notably, the absence of toxicity in blood biochemistry and histopathology of major organs after high-dose administration echoes international studies confirming lycopene’s safety in rodent models[35]. The absence of hepatorenal toxicity at therapeutic doses addresses critical safety concerns raised in oral administration studies [36], suggesting localized delivery may overcome systemic bioavailability limitations. However, our work innovatively extends these findings by establishing a direct dose-response relationship in lung cancer models and confirming the feasibility of localized intratumoral delivery—a strategy less explored compared to oral or intraperitoneal routes in previous research.

The in vitro experiments using A549 and LLC cells revealed that lycopene (10, 20, 40 μM) inhibited proliferation, clonogenicity, and promoted apoptosis, aligning with its reported effects on cell cycle arrest in colon cancer [37]. This dual-model approach enhances biological relevance given the distinct genetic backgrounds of these cell lines (adenocarcinoma vs. squamous carcinoma). Crucially, our study provides the first evidence linking lycopene’s anti-tumor effects to PI3K/AKT pathway inhibition in lung cancer, both in vivo and in vitro experiments have been verified. The observed downregulation of p-PI3K and subsequent apoptosis rescue experiments (via PI3K silencing or overexpression) established causal relationships between pathway modulation and therapeutic outcomes, resolving ambiguities in correlative studies from earlier research. This mechanistic discovery contrasts with prior studies emphasizing lycopene’s antioxidant properties, suggesting tissue-specific pathway modulation. The 40% tumor volume reduction at maximal dose exceeds reported efficacy of intraperitoneal administration in breast cancer models [38, 39], potentially attributable to optimized local bioavailability. Furthermore, the identification of PI3K/AKT axis as a primary target contrasts with proposed STAT3-mediated mechanisms in glioma [40, 41], suggesting tissue-specific pathway regulation. The innovation of this study is to systematically reveal for the first time the dual regulatory role of lycopene on the PI3K/AKT pathway in lung cancer: inhibition of pro-survival signaling and activation of mitochondria-dependent apoptotic pathways

The tumor-specific efficacy combined with systemic safety suggests lycopene’s potential as an adjuvant therapy. However, limitations exist: 1) Intratumoral delivery, while effective in mice, requires optimization for human applications; 2) Potential cross-talk with other pathways (e.g., MAPK/ERK) warrants further exploration.

## Conclusion

This study advances our understanding of lycopene’s anti-lung cancer mechanisms through dual-model validation, establishing PI3K/AKT inhibition as a key pathway and providing critical preclinical evidence for localized administration strategies. Future research should focus on nanoparticle-based delivery systems and combinatorial therapies to enhance translational potential.

## Author Contributions

Haibo Gu and Jian Ye designed the study. Chengyu Pan, Qi Xu provided the databases. Jingjing Lu and Tian Zhao assembled and analyzed the data. Keyan Fu, Xianting Yan and Yun Xu wrote the manuscript. The final submitted version has been approved by all authors.

## Ethics Statement

All experimental mice were housed in SPF-grade animal facilities at Zhejiang University’s Experimental Animal Center. All animal experiments were approved by the Zhejiang University Animal Welfare and Ethics Review Committee (ZJU20250155). CO₂ asphyxiation was used at a fill rate of 30-70% chamber volume/min, followed by cervical dislocation as a secondary physical method. All efforts were made to minimize suffering.

